# *Pseudomonas aeruginosa* increases the sensitivity of biofilm-grown *Staphylococcus aureus* to membrane-targeting antiseptics and antibiotics

**DOI:** 10.1101/668780

**Authors:** Giulia Orazi, Kathryn L. Ruoff, George A. O’Toole

**Affiliations:** Department of Microbiology and Immunology, Geisel School of Medicine at Dartmouth, Hanover, New Hampshire, USA

**Keywords:** *Pseudomonas aeruginosa*, *Staphylococcus aureus*, membrane, antibiotics, biofilm

## Abstract

*Pseudomonas aeruginosa* and *Staphylococcus aureus* often cause chronic, recalcitrant infections in large part due to their ability to form biofilms. The biofilm mode of growth enables these organisms to withstand antibacterial insults that would effectively eliminate their planktonic counterparts. We found that *P. aeruginosa* supernatant increased the sensitivity of *S. aureus* biofilms to multiple antimicrobial compounds, including fluoroquinolones and membrane-targeting antibacterial agents, including the antiseptic chloroxylenol. Treatment of *S. aureus* with the antiseptic chloroxylenol alone did not decrease biofilm cell viability; however, the combination of chloroxylenol and *P. aeruginosa* supernatant led to a 4-log reduction in *S. aureus* biofilm viability compared to exposure to chloroxylenol alone. We found that the *P. aeruginosa*-produced small molecule 2-n-heptyl-4-hydroxyquinoline N-oxide (HQNO) is responsible for the observed heightened sensitivity of *S. aureus* to chloroxylenol. Similarly, HQNO increased the susceptibility of *S. aureus* biofilms to other compounds, including both traditional and non-traditional antibiotics, which permeabilize bacterial membranes. Genetic and phenotypic studies support a model whereby HQNO causes an increase in *S. aureus* membrane fluidity, thereby improving the efficacy of membrane-targeting antiseptics and antibiotics. Importantly, our data show that *P. aeruginosa* exoproducts can enhance the ability of various antimicrobial agents to kill biofilm populations of *S. aureus* that are typically difficult to eradicate, providing a path for the discovery of new biofilm-targeting antimicrobial strategies.

**Importance:** The thick mucus in the airways of cystic fibrosis (CF) patients predisposes them to frequent, polymicrobial respiratory infections. *Pseudomonas aeruginosa* and *Staphylococcus aureus* are frequently co-isolated from the airways of individuals with CF, as well as from diabetic foot ulcers and other wounds. Both organisms form biofilms, which are notoriously difficult to eradicate and promote chronic infection. In this study, we have shown *P. aeruginosa* secreted factors can increase the efficacy of compounds that alone have little or no bactericidal activity against *S. aureus* biofilms. In particular, we discovered that *P. aeruginosa* exoproducts can potentiate the anti-staphylococcal activity of phenol-based antiseptics and other membrane-active drugs, including non-traditional antibiotics. Our findings illustrate that polymicrobial interactions can dramatically increase antibacterial efficacy *in vitro*, and may guide new approaches to target persistent infections, such as those commonly found in respiratory tract infections and in chronic wounds.

## Introduction

Bacterial biofilms are the underlying cause of many chronic, difficult-to-treat infections. The biofilm lifestyle confers high-level tolerance to antibiotics and antiseptics, which is reflected by the requirement of 100-1000 times higher concentrations of these compounds to treat biofilms compared to their planktonic counterparts (1). As a result, it has proven difficult to find treatments that effectively eradicate biofilms (2-4).

Studies assessing biofilm antibiotic and antiseptic tolerance have typically been performed with single-species biofilms. While such single-species communities are commonly associated with implant infections (5), many infections are caused by polymicrobial biofilms, including respiratory infections, otitis media, urinary tract infections, and infections of both surgical and chronic wounds (6-19). Emerging evidence suggests that growth in these mixed microbial communities can alter antimicrobial tolerance profiles, often in unexpected ways (20-40), but the mechanism(s) underlying such altered tolerance are often poorly understood, with some exceptions. For example, a previous study from our group showed that secreted products of *Pseudomonas aeruginosa* could enhance biofilm tolerance of *Staphylococcus aureus* to vancomycin by 100-fold, likely via interfering with the function of the electron transport chain and slowing growth of *S. aureus* (37).

*P. aeruginosa* and *S. aureus* coexist in multiple infection settings, and both form biofilms that can be difficult to eradicate. *P. aeruginosa* and *S. aureus* are two of the most prevalent respiratory pathogens in patients with cystic fibrosis (CF), and are both associated with poor lung function and clinical outcomes in these patients (41-45). CF patients who are co-infected with *P. aeruginosa* and *S. aureus* have worse outcomes than those who are infected with either organism alone (46-50). In addition, *P. aeruginosa* and *S. aureus* are often co-isolated from chronic wounds, including difficult-to-treat diabetic foot ulcers (51, 52). Furthermore, *in vitro* evidence suggests that *P. aeruginosa* and *S. aureus* coinfection delays wound healing (53).

In this study, we have identified several compounds that alone have little activity against *S. aureus* biofilms, but when combined with secreted products from *P. aeruginosa*, these agents can effectively decrease *S. aureus* biofilm viability. We propose a model whereby the *P. aeruginosa* exoproduct 2-n-heptyl-4-hydroxyquinoline N-oxide (HQNO) interacts with the *S. aureus* cell membrane, which leads to increased membrane fluidity and potentiates the ability of membrane-active compounds to more effectively target *S. aureus* biofilms.

## Results

### *P. aeruginosa* supernatant increases *S. aureus* sensitivity to multiple antibiotic compounds

In a previous study, we found that *P. aeruginosa* exoproducts decrease the efficacy of vancomycin against *S. aureus* biofilms (37). To test whether *P. aeruginosa* might impact *S. aureus* sensitivity to other antibiotics, we screened Biolog Phenotype MicroArray Panels for changes in *S. aureus* antibiotic sensitivity in the presence versus absence of *P. aeruginosa* cell-free culture supernatant. We identified many compounds that became either less effective, as reported previously (37), or as we show here, more effective at killing *S. aureus* when in the presence of *P. aeruginosa* exoproducts (Table S1).

Among the several classes of antimicrobial agents that became more effective at killing *S. aureus* in the presence of *P. aeruginosa* supernatant are nucleic acid synthesis inhibitors, membrane-active antibiotics, and antiseptics. Additionally, we identified other compounds that are not typically used to treat bacterial infections that became more effective at decreasing *S. aureus* viability, including anti-cholinergic agents, antipsychotic drugs, and ion channel blockers (Table S1).

### *P. aeruginosa* supernatant increases *S. aureus* biofilm sensitivity to chloroxylenol

In the experiments described above using the Biolog Phenotype MicroArray Panels, the compounds tested were added at the same time as the microbes were inoculated into the medium, thus there was limited time for the bacteria to form a biofilm before exposure to the candidate agents. Therefore, we next tested whether *P. aeruginosa* supernatant could increase the efficacy of the compounds we identified in the Biolog screen against pre-formed *S. aureus* biofilms. In these experiments, the biofilm of *S. aureus* Newman was allowed to form for 6 h, and fresh medium supplemented with the indicated compound and/or *P. aeruginosa* supernatant was added to this preformed biofilm. This method is what we refer to as the *biofilm disruption assay*, described in more detail in the Supplemental Materials and Methods (Text S1). Previously, we showed that by 6 h post-inoculation (p.i.) the adherent population of *S. aureus* Newman cells is tolerant to vancomycin; at this time point, there is a difference of 3 logs between the cell viability of the biofilm population compared to the planktonic population for a given dose of antibiotic (37). Thus, these communities have one of the key phenotypic traits of a biofilm.

We found that *P. aeruginosa* supernatant increased the sensitivity of early (6 h) *S. aureus* biofilms to the topical antibiotic chloroxylenol (Fig. 1). Similar to other phenol-based antiseptics, this compound impacts bacterial cell membranes, leading to increased fluidity and membrane permeability (54-56). Alone, chloroxylenol displayed modest activity against *S. aureus* biofilms. Strikingly, the ability of the antiseptic chloroxylenol to kill early *S. aureus* Newman biofilms was enhanced by 4 logs compared to the activity of chloroxylenol alone when combined with *P. aeruginosa*-secreted products (Fig. 1). We evaluated whether this phenotype is specific to the Newman strain or a more general phenomenon by testing multiple *S. aureus* laboratory strains and clinical isolates – both methicillin-sensitive and methicillin-resistant (Table S2). In all cases, we observed that *P. aeruginosa* supernatant dramatically increased the efficacy of chloroxylenol against *S. aureus* biofilms (Fig. 1). Chloroxylenol is dissolved in ethanol; we confirmed that the volume of ethanol used does not decrease *S. aureus* viability in either the presence or absence of *P. aeruginosa* supernatant (Fig. S1A). Moreover, the impact of supernatant on *S. aureus* sensitivity to chloroxylenol could be observed as early as 3 h after addition of the compounds to a 6 h old biofilm, and the reduction in viability continued for 24 h post-treatment, wherein the assay was reaching its limit of detection (Fig. S1B).

**Figure 1.**
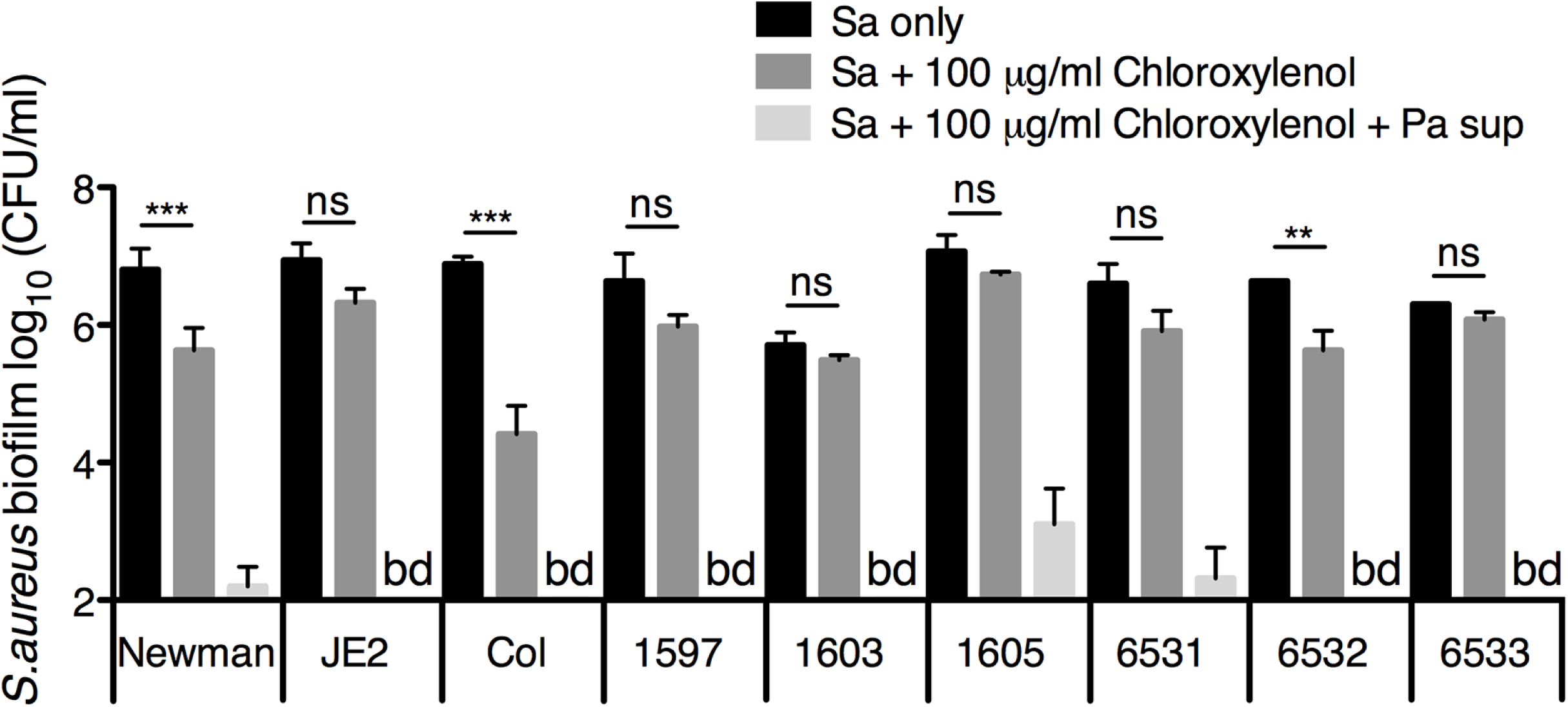
*P. aeruginosa* supernatant increases *S. aureus* biofilm sensitivity to chloroxylenol. Biofilm disruption assays on plastic were performed with the specified *S. aureus* clinical isolate, *P. aeruginosa* PA14 supernatant (Pa sup), and chloroxylenol (Chlor) at 100 μg/ml. Biofilms were grown for 6 hours, exposed to the above treatments for 18 hours, and *S. aureus* biofilm CFU were determined. Each column displays the average from two biological replicates, each with three technical replicates. Error bars indicate standard deviation (SD). bd, below detection. ns, not significant; **, *P* < 0.01, ***, *P* < 0.001, by ordinary one-way ANOVA and Bonferroni’s multiple comparison post-test.

### *P. aeruginosa* supernatant increases the ability of chloroxylenol to eradicate difficult-to-treat *S. aureus* biofilms

We then determined whether *P. aeruginosa* could enable chloroxylenol to kill especially difficult-to-treat *S. aureus* biofilms. *S. aureus* grown in anoxia and respiration-deficient *S. aureus* small colony variants (SCVs) both exhibit high tolerance to many classes of antibiotics (57-59), likely because the bacteria need to be actively growing in order for many antibacterial compounds to be effective. Depending on the antibiotic class, either the antibiotic target needs to be produced or electron transport is required for drug uptake (57, 60), but membrane-targeting agents are an exception; the target is present whether or not the organism is actively growing (61). Indeed, *P. aeruginosa* supernatant increased the efficacy of chloroxylenol against *S. aureus* Newman biofilms to similar degrees in anoxia and normoxia (Fig. 2A).

**Figure 2.**
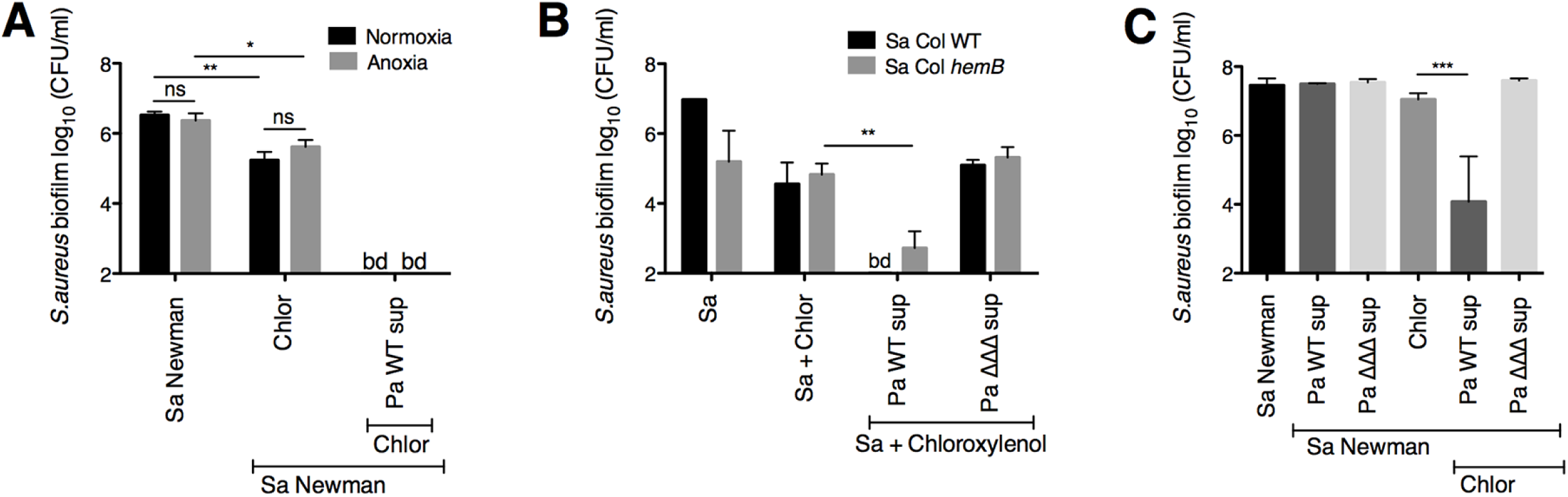
*P. aeruginosa* supernatant enhances the ability of chloroxylenol to kill difficult-to-treat *S. aureus* biofilms. (**A**) Biofilm disruption assays on plastic were performed with *S. aureus* (Sa) Newman, *P. aeruginosa* PA14 supernatant (Pa sup), and chloroxylenol (Chlor) at 100 μg/ml under normoxic or anoxic conditions. Biofilms were grown for 6 hours, exposed to the above treatments for 18 hours, and *S. aureus* biofilm CFU were determined. **(B)** Biofilm disruption assays on plastic were performed with *S. aureus* (Sa) Col parental strain or *hemB* mutant, supernatants from wild-type *P. aeruginosa* PA14 and the Δ*pqsL* Δ*pvdA* Δ*pchE* mutant (Pa ΔΔΔ sup), and chloroxylenol (Chlor) at 100 μg/ml. Biofilms were grown for 6 hours, exposed to the above treatments for 18 hours, and *S. aureus* biofilm CFU were determined. (**C**) Biofilm disruption assays on plastic were performed with *S. aureus* (Sa) Newman, supernatants from wild-type *P. aeruginosa* PA14 and the Δ*pqsL* Δ*pvdA* Δ*pchE* mutant (Pa ΔΔΔ sup), and chloroxylenol (Chlor) at 100 μg/ml. Biofilms were grown for 24 hours, exposed to the above treatments for 24 additional hours, and *S. aureus* biofilm CFU were determined. Each column displays the average from three biological replicates, each with three technical replicates. Error bars indicate SD. bd, below detection. ns, not significant; *, *P* < 0.05, **, *P* < 0.01, ***, *P* < 0.001, by ordinary one-way ANOVA and Tukey’s multiple comparison post-test.

To test whether the combination of *P. aeruginosa* supernatant and chloroxylenol is effective against biofilm-grown *S. aureus* SCVs, we used a *S. aureus* Col strain that has a mutation in *hemB*, a gene involved in hemin biosynthesis. The *S. aureus hemB* mutant is defective in electron transport and has the typical characteristics of clinical SCVs (62). We observed that *P. aeruginosa* supernatant enhanced chloroxylenol’s activity against the Col *hemB* mutant as well as the parental strain (Fig. 2B).

Furthermore, we tested whether more mature *S. aureus* biofilms could be effectively targeted by the *P. aeruginosa* supernatant-chloroxylenol combination. When we grew *S. aureus* Newman biofilms for 24 h before exposure to the combination treatment, we observed a striking 4 log-fold enhancement of chloroxylenol’s antimicrobial activity (Fig. 2C), similar to what was seen for 6 h-grown biofilms (Fig. 1).

### The *P. aeruginosa* exoproducts HQNO and siderophores increase *S. aureus* biofilm and planktonic sensitivity to chloroxylenol

To explore the mechanism underlying *P. aeruginosa* supernatant-mediated enhancement of chloroxylenol’s anti-staphylococcal activity, we sought to identify *P. aeruginosa* mutants that were unable to increase the sensitivity of *S. aureus* Newman biofilms to this drug. Previously, we showed that 2-n-heptyl-4-hydroxyquinoline N-oxide (HQNO) and siderophores contribute to the ability of *P. aeruginosa*-to protect *S. aureus* from vancomycin (37). Thus, we tested *P. aeruginosa* PA14 strains with mutations in genes encoding components of the *Pseudomonas* quinolone signal (PQS) quorum sensing system (*pqsA, pqsH, pqsL*), and biosynthesis of the siderophores pyoverdine (*pvdA*) and pyochelin (*pchE*). Supernatants from *P. aeruginosa* PA14 Δ*pqsA*, Δ*pqsH*, Δ*pqsL*, and Δ*pvdA* Δ*pchE* mutants each had a defect in the ability to increase *S. aureus* Newman biofilm sensitivity to chloroxylenol relative to the wild-type *P. aeruginosa* PA14 (Fig. S2A, B).

Additionally, we tested *P. aeruginosa* PA14 strains with mutations in genes encoding the following secreted products: hydrogen cyanide (*hcnA, hcnB*), LasA protease (*lasA*), elastase (*lasB*), and rhamnolipids (*rhlA*). Supernatants from these mutants retained the ability to increase the sensitivity of *S. aureus* biofilms to chloroxylenol (Fig. S2A, B).

To investigate whether HQNO, pyoverdine, and pyochelin all contributed to the phenotype, we tested whether the supernatant from *P. aeruginosa* strains with mutations in the genes encoding all three factors was deficient in enhancing chloroxylenol’s activity against *S. aureus*. Indeed, supernatant from the *P. aeruginosa* PA14 Δ*pqsL* Δ*pvdA* Δ*pchE* mutant (designated the ΔΔΔ mutant) was unable to increase the sensitivity of *S. aureus* Newman biofilms to chloroxylenol (Fig. S2B, 3A). Supernatant from the *P. aeruginosa* PA14 Δ*pqsL* Δ*pvdA* Δ*pchE* mutant was unable to potentiate the ability of chloroxylenol to kill difficult-to-treat SCVs and 24 h-grown biofilms (Fig. 2B, C; Pa ΔΔΔ sup). Similar to the biofilm population, we observed that *P. aeruginosa* PA14 wild-type supernatant, but not the Δ*pqsL* Δ*pvdA* Δ*pchE* mutant, enhances the ability of chloroxylenol to kill planktonic *S. aureus* Newman by approximately 3 logs (Fig. 3B). Thus, our data indicate that HQNO and both siderophores are required for *P. aeruginosa*-mediated enhancement of chloroxylenol’s activity against both planktonic and biofilm populations of *S. aureus*.

**Figure 3.**
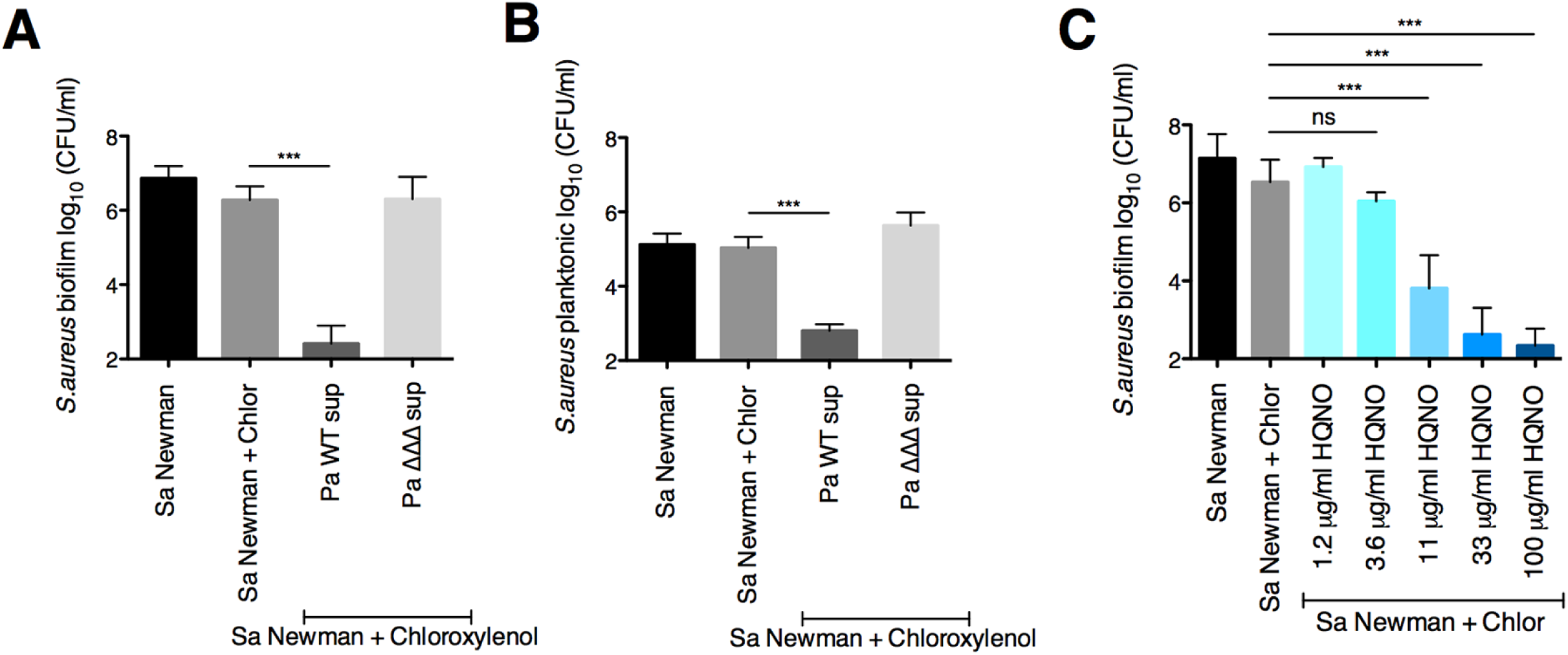
The *P. aeruginosa* exoproducts HQNO and siderophores increase *S. aureus* biofilm and planktonic sensitivity to chloroxylenol. (**A and B**) Biofilm disruption assays on plastic were performed with *S. aureus* (Sa) Newman, supernatants from *P. aeruginosa* PA14 wild-type and the Δ*pqsL* Δ*pvdA* Δ*pchE* deletion mutant (Pa ΔΔΔ sup), and chloroxylenol (Chlor) at 100 μg/ml. Biofilms were grown for 6 hours, exposed to the above treatments for 18 hours, and *S. aureus* biofilm (**A**) and planktonic (**B**) CFU were determined. Data in panels A and B were from the same experiments. (**C**) Biofilm disruption assays on plastic were performed with *S. aureus* (Sa) Newman, chloroxylenol (Chlor) at 100 μg/ml, and the specified concentrations of HQNO (dissolved in DMSO). Biofilms were grown for 6 hours, exposed to the above treatments for 18 hours, and *S. aureus* biofilm CFU were determined. Each column displays the average from at least three biological replicates, each with three technical replicates. Error bars indicate SD. ns, not significant; ***, *P* < 0.001, by ordinary one-way ANOVA and Tukey’s multiple comparison post-test.

### HQNO alone enhances the activity of chloroxylenol against *S. aureus* biofilms

To test whether HQNO alone could enhance to the ability of chloroxylenol to kill *S. aureus* in biofilm, we performed a biofilm disruption assay using commercially available HQNO. We used concentrations of HQNO that are in the range of those produced by *P. aeruginosa* PA14 under our experimental conditions (37), as well as those produced by stationary-phase *P. aeruginosa* cultures grown in rich media (63, 64). Previously, we quantified the level of HQNO produced by *P. aeruginosa* PA14 after 24 h of growth in minimal medium on plastic plates, which is the source of *P. aeruginosa* supernatants used throughout this study (37). We found that the level of HQNO in these *P. aeruginosa* supernatants is ∼10 μg/ml. Additionally, *P. aeruginosa* PA14 produced ∼15 μg/ml HQNO when grown on CF-derived epithelial cells for 6 h (37). We observed a dose-response whereby increasing concentrations of exogenous HQNO corresponded with enhanced ability of chloroxylenol to kill *S. aureus* Newman biofilms (Fig. 3C). These results indicate that the presence of a single secreted factor, HQNO, is sufficient to alter *S. aureus* biofilm sensitivity to chloroxylenol.

### HQNO likely does not increase *S. aureus* sensitivity to chloroxylenol via inhibition of the electron transport chain

HQNO is well known to inhibit Complex II and III of the *S. aureus* electron transport chain (ETC) (65-68). To investigate whether HQNO shifts *S. aureus* sensitivity to chloroxylenol by inhibiting respiration, we tested the following ETC inhibitors: 3-Nitropropionic acid (3-NP; Complex II inhibitor), Antimycin A (Complex III inhibitor), sodium azide (azide; Complex IV inhibitor), and Oligomycin (ATP synthase inhibitor) or mutations in components of ATP synthase. All but one of the compounds tested, Antimycin A, had little to no impact on *S. aureus* sensitivity to chloroxylenol, nor did mutations in the ATPase (Fig. S3A-E).

It is possible that HQNO and Antimycin A are changing antibiotic sensitivity not by inhibiting the ETC, but via a different mechanism entirely. Thus, we took a different approach to investigate whether ETC inhibition changes *S. aureus* susceptibility to chloroxylenol. Exposure to anoxic conditions is a way to inhibit respiration that does not require the use of chemical compounds. Anoxia did not enhance chloroxylenol’s efficacy against *S. aureus* Newman biofilms in the absence of *P. aeruginosa* supernatant (Fig. 2A). Also, despite lacking a functional ETC, *S. aureus* SCVs are not hypersensitive to chloroxylenol (Fig. 2B). Furthermore, as we observed above, *P. aeruginosa* supernatant is able to potentiate the activity of chloroxylenol to kill SCVs even though these cells are respiration-deficient (Fig. 2B). Together, these data indicate that HQNO likely alters *S. aureus* antibiotic sensitivity via a mechanism independent of its effects on the ETC.

We next considered several possible mechanisms underlying HQNO-mediated enhancement of chloroxylenol’s anti-staphylococcal activity. Specifically, we tested the following models: 1) HQNO-mediated changes in membrane potential increase antibiotic sensitivity, 2) HQNO-induced generation of reactive oxygen species leads to enhanced bacterial killing, 3) HQNO alters the ability of *S. aureus* to efflux chloroxylenol, and/or 4) HQNO changes properties of the *S. aureus* cell membrane. Experiments testing the first three of these models, which did not support these models, are presented in the Supplemental Results (Text S1) and in Figures S3-S4.

### Exogenous HQNO increases *S. a ureus* membrane fluidity

Previous studies have found that changes in the cell membrane fatty acid composition, which influences membrane fluidity, alter the susceptibility of bacterial cells to phenolic compounds (69). Thus, we tested whether HQNO might cause heightened susceptibility to chloroxylenol by altering the fluidity of the *S. aureus* cell membrane. To measure membrane fluidity, we performed Laurdan generalized polarization (GP) assays. Laurdan is a fluorescent dye that is sensitive to changes in membrane fluidity; the emission spectrum changes depending on the physical state of lipids within a bilayer. A decrease in Laurdan GP values corresponds to an increase in membrane fluidity. This dye has been previously used to measure the cell membrane fluidity of *S. aureus* (70-72).

We used benzyl alcohol, a well-established membrane fluidizing agent (73-75), as a positive control. Exposure to 500 mM or 1 M of benzyl alcohol for 1 h led to a significant decrease in Laurdan GP relative to *S. aureus* exposed to MEM, indicating an increase in membrane fluidity (Fig. 4A). We observed that treatment of *S. aureus* Newman with HQNO at all concentrations tested led to a significant reduction in Laurdan GP relative to exposure to MEM alone, indicating that HQNO has a fluidizing effect on the *S. aureus* membrane (Fig. 4B). Additionally, we found that exposure to Antimycin A also led to a significant increase in fluidity (Fig. 4C), albeit to a lesser extent than HQNO (Fig. 4B). Furthermore, we showed that the solvents for HQNO and Antimycin A, DMSO and ethanol, respectively, did not cause the observed increase in *S. aureus* membrane fluidity (Fig. 4B, C).

**Figure 4.**
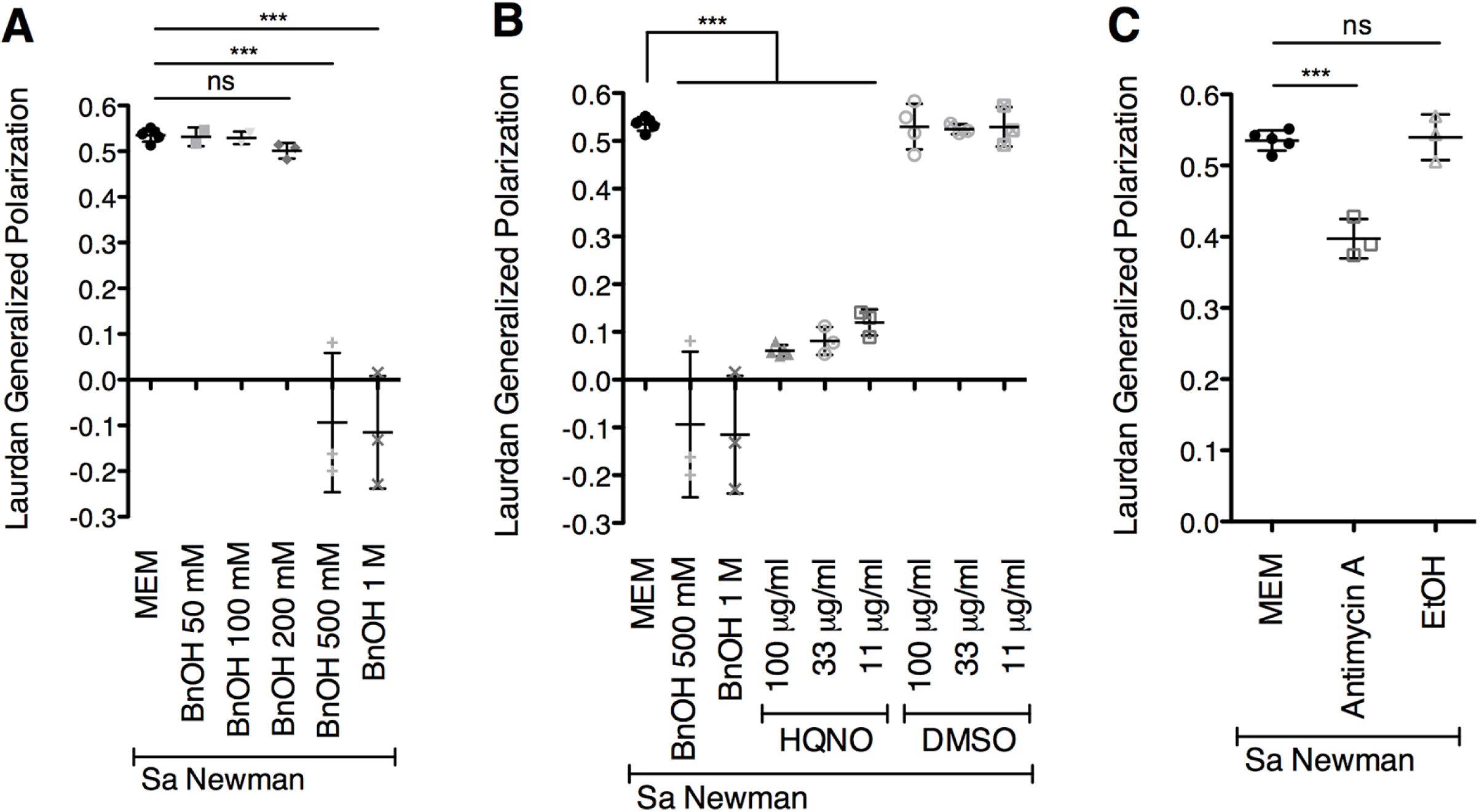
Exogenous HQNO increases *S. aureus* membrane fluidity. **(A to C)** Laurdan generalized polarization (GP) was performed with *S. aureus* (Sa) Newman, benzyl alcohol (BnOH) (**A and B**), HQNO (**B**), and the DMSO control (solvent for HQNO) at the indicated concentrations, and Antimycin A at 100 μg/ml along with the ethanol control (solvent for Antimycin A) (**C**). *S. aureus* was exposed to the above treatments for 1 hour, and GP values were determined. Each column displays the average from at least three biological replicates, each with three technical replicates. Error bars indicate SD. ns, not significant; ***, *P* < 0.001, by ordinary one-way ANOVA and Tukey’s multiple comparison post-test.

### Shifting membrane fluidity alters *S. aureus* biofilm sensitivity to chloroxylenol

Next, we investigated whether the observed HQNO-mediated increase in membrane fluidity can lead to increased sensitivity to chloroxylenol. To test this hypothesis, we exposed *S. aureus* biofilms to various compounds that are known to influence membrane fluidity. Benzyl alcohol and 1-heptanol both impart higher fluidity, whereas dimethyl sulfoxide (DMSO) causes membranes to become less fluid (73-77). We observed that benzyl alcohol and 1-heptanol both increase *S. aureus* Newman biofilm sensitivity to chloroxylenol (Fig. 5A, B). In contrast, the membrane-rigidifying agent DMSO did not increase *S. aureus* Newman biofilm sensitivity to chloroxylenol (Fig. 5C). These results suggest that alterations in *S. aureus* membrane fluidity impact sensitivity to chloroxylenol, whereby increased fluidity leads to higher sensitivity.

**Figure 5.**
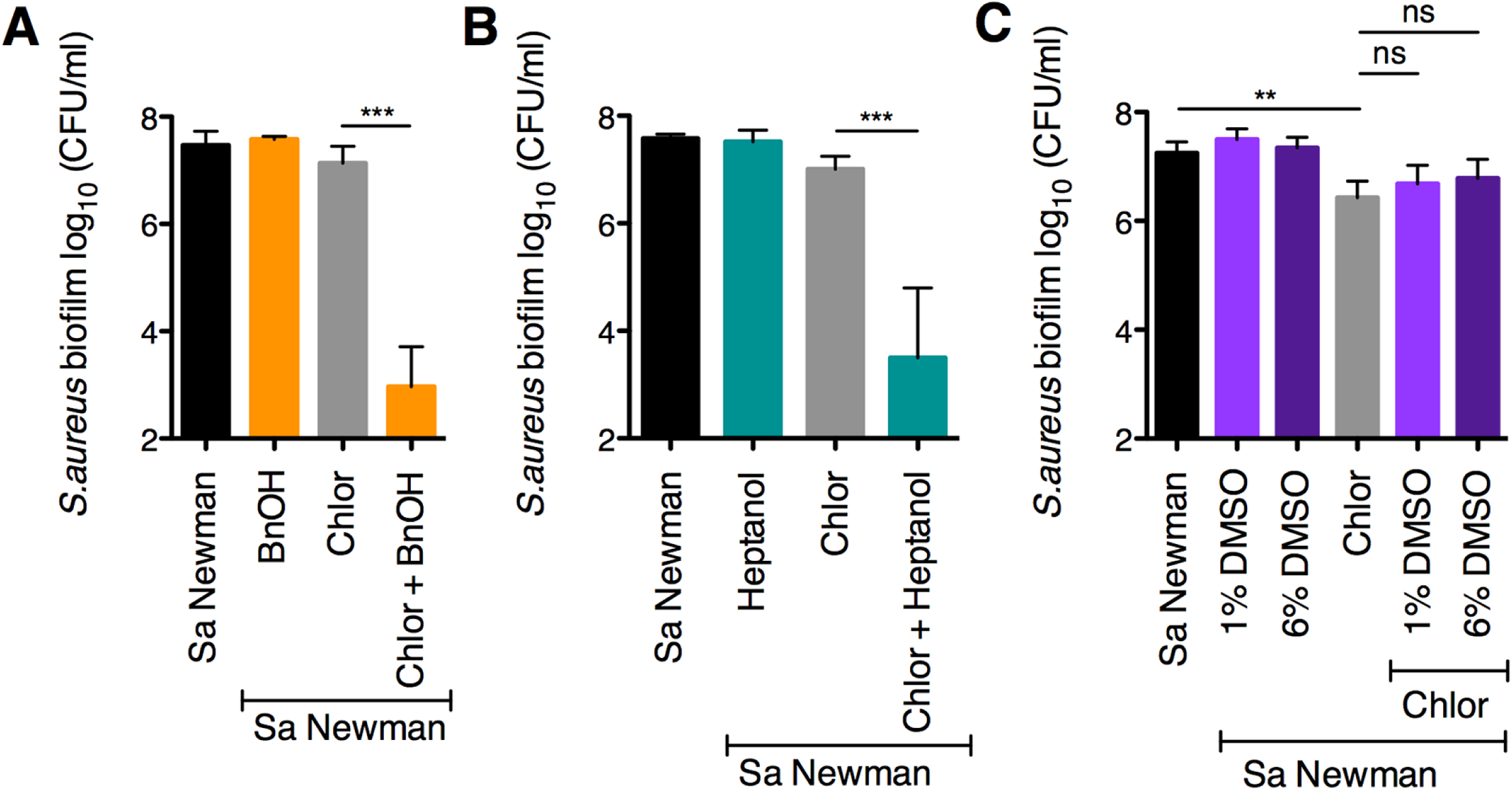
Shifting membrane fluidity alters *S. aureus* biofilm sensitivity to chloroxylenol. (**A to C**) Biofilm disruption assays on plastic were performed with *S. aureus* (Sa) Newman, chloroxylenol (Chlor) at 100 μg/ml, benzyl alcohol (BnOH) at 50 mM (**A**), 1-heptanol at 50 mM (**B**), and dimethyl sulfoxide (DMSO) at 1% and 6% (**C**). Biofilms were grown for 6 hours, exposed to the above treatments for 18 hours, and *S. aureus* biofilm CFU were determined. Each column displays the average from at least three biological replicates, each with three technical replicates. Error bars indicate SD. ns, not significant; **, *P* < 0.01, ***, *P* < 0.001, by ordinary one-way ANOVA and Tukey’s multiple comparison post-test.

Next, we showed that manipulating *S. aureus* fatty acid composition either by adding exogenous unsaturated fatty acids (Fig S5A, Text S1) or increasing the proportion of branched-chain fatty acids (BCFAs) relative to short chain fatty acids by mutation (SCFAs; Fig. S5B, Text S1) leads to increased *S. aureus* sensitivity to chloroxylenol. Additionally, we showed that decreasing levels of BCFAs relative to SCFAs by introducing the *lpd* mutation does not increase sensitivity to chloroxylenol (Fig. S5C), and that cardiolipin is not required for altered *S. aureus* sensitivity to this drug (Fig. S5D, Text S1).

Together, our data suggest that changes in membrane fatty acid composition influence the efficacy of chloroxylenol and are consistent with our model that an increase in membrane fluidity promotes chloroxylenol’s ability to kill *S. aureus* biofilms.

### Prolonged exposure to *P. aeruginosa* exoproducts alters *S. aureus* membrane fatty acid profiles

Our data above suggest that HQNO increases *S. aureus* membrane fluidity, which leads to heightened sensitivity of *S. aureus* to chloroxylenol. Thus, we explored whether HQNO induces changes in *S. aureus* membrane fatty acid composition. We performed a time course to track *S. aureus* fatty acid composition over time in the presence of *P. aeruginosa* exoproducts. Briefly, *S. aureus* Newman cells were exposed to medium alone (MEM + L-Gln) or *P. aeruginosa* PA14 wild-type supernatant for differing lengths of time (30 min, 1 h, 3 h, 6 h, or 10 h). Subsequently, fatty acid methyl ester (FAME) analysis was performed to measure the membrane fatty acid composition.

By 30 min or 1 h, the membrane fatty acid profile of *S. aureus* cells grown in medium alone appeared similar to the profile of *P. aeruginosa* supernatant-exposed *S. aureus* cells (Fig. S6A-C, Table S3). However, prolonged treatment with *P. aeruginosa* supernatant led to a shift in *S. aureus* membrane fatty acid profiles. In particular, *S. aureus* cells incubated with *P. aeruginosa* exoproducts for 24 h had significantly reduced relative BCFA levels compared to *S. aureus* grown in medium alone (Fig. S6D-E, Text S1). Above, we found that HQNO significantly increases *S. aureus* membrane fluidity after 1 h (Fig. 4B). Because the fluidizing effect of HQNO occurs more rapidly compared to the effect of *P. aeruginosa* supernatant on *S. aureus* membrane fatty acid composition, it is likely that the HQNO-mediated increase in *S. aureus* membrane fluidity we observe does not occur via changes in membrane fatty acid profiles.

### *P. aeruginosa* supernatant increases *S. aureus* biofilm sensitivity to multiple membrane-targeting compounds

Given the effects of *P. aeruginosa* exoproducts on *S. aureus* sensitivity to chloroxylenol, we explored whether *P. aeruginosa* alters the anti-staphylococcal efficacy of other membrane-active antibiotics. Here, we tested the efficacy of the phenol-based antiseptic biphenyl, as well as the topical peptide antibiotic gramicidin in combination with *P. aeruginosa* supernatant. Both of these compounds are thought to kill bacteria by ultimately causing an increase in cell membrane permeability. We discovered that *P. aeruginosa* secreted products enhance the ability of the membrane-active drugs biphenyl and gramicidin to kill *S. aureus* Newman biofilms (Fig. 6A, B). We also made the interesting observation that *P. aeruginosa* supernatant increases *S. aureus* biofilm sensitivity to two non-traditional antibiotics, trifluoperazine, an antipsychotic, and amitriptyline, an antidepressant (Fig. 6C, D). Strikingly, the combination of either of these drugs and *P. aeruginosa* supernatant led to a 2.5 to 3-log reduction in *S. aureus* biofilm viability compared to exposure to the drug alone (Fig. 6C, D). Supernatants from *P. aeruginosa* PA14 Δ*pqsL* and Δ*pvdA* Δ*pchE* mutants each had defects in the ability to increase *S. aureus* Newman biofilm sensitivity to trifluoperazine and amitriptyline relative to the wild-type *P. aeruginosa* PA14 (Fig. 6C, D), suggesting that HQNO and siderophores both contribute to this phenotype. In contrast, it appears than another *P. aeruginosa*-produced factor is involved in enhancing the activity of gramicidin against *S. aureus* biofilms (Fig. 6B).

**Figure 6.**
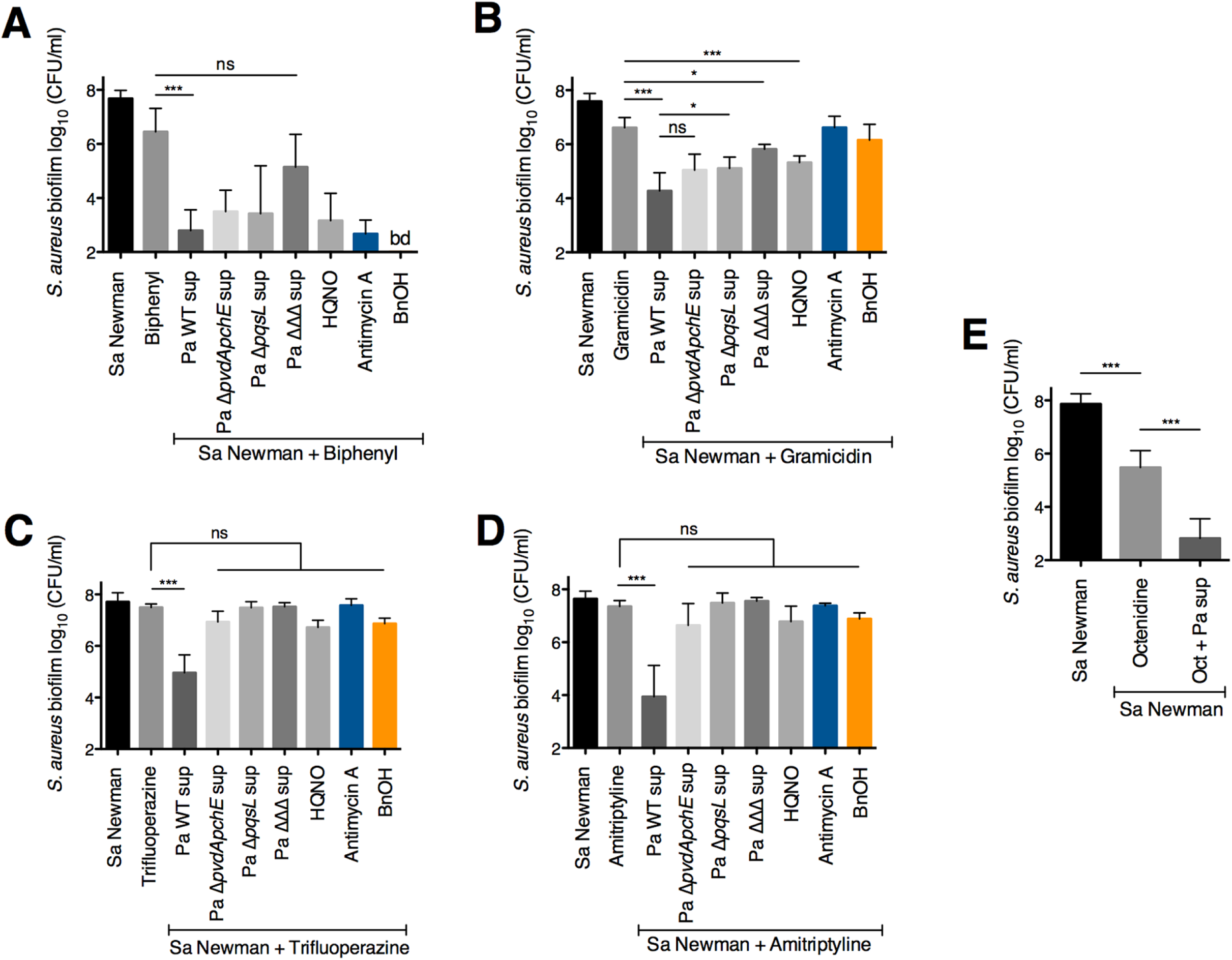
*P. aeruginosa* supernatant increases *S. aureus* biofilm sensitivity to other membrane-targeting compounds. (**A to E**) Biofilm disruption assays on plastic were performed with *S. aureus* (Sa) Newman, supernatants from *P. aeruginosa* PA14 wild-type and the specified mutants (Pa sup), and either Biphenyl at 200 μg/ml (**A**), Gramicidin at 100 μg/ml (**B**), Trifluoperazine at 100 μg/ml (**C**), Amitriptyline at 100 μg/ml (**D**), or Octenidine dihydrochloride (Oct) at 5 μg/ml (**E**). Biofilms were grown for 6 hours, exposed to the above treatments for 18 hours, and *S. aureus* biofilm CFU were determined. Each column displays the average from at least three biological replicates, each with three technical replicates. Error bars indicate standard deviation (SD). ns, not significant; *, *P* < 0.05, ***, *P* < 0.001, by ordinary one-way ANOVA and Tukey’s multiple comparison post-test.

Additionally, we examined whether altering membrane fluidity influenced *S. aureus* biofilm sensitivity to the above compounds. We observed that benzyl alcohol did not appreciably alter *S. aureus* sensitivity to gramicidin, trifluoperazine, or amitriptyline (Fig. 6B-D). In contrast, the fluidizing agent led to a striking increase in the antibacterial efficacy of biphenyl; the combination of these compounds led to a decrease in *S. aureus* Newman biofilm viability to below the level of detection of this assay (∼200 CFU/ml, Fig. 6A). These results suggest that a more fluid membrane increases the susceptibility of *S. aureus* biofilms to biphenyl, which is a compound similar to chloroxylenol in structure and function.

Finally, we tested whether *P. aeruginosa* secreted products could increase the anti-staphylococcal efficacy of octenidine dihydrochloride, a surfactant-based antiseptic that is approved for treatment of wound infections and has low cytotoxicity (78, 79). We observed that *P. aeruginosa* supernatant potentiates the activity of octenidine against *S. aureus* biofilms by 2.5 logs (Fig. 6E).

## Discussion

In this study, we found that the interactions between two bacterial pathogens that are frequently co-isolated from infections can cause striking and unexpected changes in antimicrobial susceptibility profiles. We showed that *P. aeruginosa* potentiates the ability of various antibacterial agents to kill *S. aureus* biofilms, which are often difficult to eradicate. In particular, we found that *P. aeruginosa* secreted products increase the sensitivity of *S. aureus* biofilms to the topical antiseptic chloroxylenol. Alone, chloroxylenol at a concentration of 100 μg/ml is not effective at eradicating *S. aureus* biofilms; however, in combination with *P. aeruginosa* cell-free culture supernatant, which alone does not impact *S. aureus* viability, the efficacy of chloroxylenol increased 4 log-fold. Moreover, we have shown that *P. aeruginosa* supernatant can increase the ability of chloroxylenol to kill multiple strains and clinical isolates of *S. aureus*. Furthermore, we found that the small molecule HQNO and the siderophores pyoverdine and pyochelin contribute to the *P. aeruginosa*-mediated increase in the efficacy of chloroxylenol against *S. aureus* biofilms. In addition, we showed that HQNO alone recapitulated the effect of *P. aeruginosa* supernatant. Thus, the addition of a small molecule alone can greatly influence the efficacy of this antiseptic.

Previous studies have detected HQNO in expectorated sputum from CF patients infected with *P. aeruginosa*, and these levels are highly variable (29, 80). *P. aeruginosa* isolates from chronic CF pulmonary infections frequently have loss-of-function mutations in the quorum sensing regulator *lasR*, and often overproduce alginate (81, 82). LasR inactivity and mucoidy each can lead to decreased HQNO production *in vitro* (64, 83). Therefore, quorum sensing activity and mucoidy may modulate the levels of HQNO produced by *P. aeruginosa* during infection, and in turn, influence the ability of HQNO to modify *S. aureus* drug sensitivity profiles *in vivo*.

HQNO has been shown to inhibit the *S. aureus* electron transport chain (ETC) (65). To investigate whether HQNO influences *S. aureus* susceptibility to chloroxylenol via inhibition of respiration, we treated *S. aureus* with chemical inhibitors of the ETC alone or in combination with the antibiotic. We found that only a subset of the ETC inhibitors tested increased the efficacy of chloroxylenol. However, anoxia did not increase *S. aureus* chloroxylenol sensitivity in the absence of HQNO. Additionally, despite having a defective ETC, *S. aureus* SCVs became more susceptible to chloroxylenol in the presence of HQNO, suggesting that inhibition of respiration is not required for this phenotype.

Since it is known that changes in membrane lipid profiles impact sensitivity to membrane-targeting compounds (69), we hypothesized that HQNO might cause heightened susceptibility to chloroxylenol by altering one or more properties of the *S. aureus* cell membrane. Like other phenol-based antiseptics, chloroxylenol is thought to insert into the cell membrane and cause an increase in membrane fluidity and permeability (54-56). Thus, an increase in membrane fluidity mediated by HQNO may allow for greater accumulation of chloroxylenol within the membrane, and subsequently cause an increase in efficacy of the antibiotic. Manipulating the fluidity of *E. coli* membranes has been previously demonstrated to alter sensitivity to phenols, whereby decreasing membrane fluidity conferred increased tolerance to these compounds (69). Therefore, we tested whether HQNO changes the fluidity of the *S. aureus* cell membrane, potentially explaining the increased antimicrobial sensitivity we observe. We found that exogenous HQNO causes a striking increase in *S. aureus* membrane fluidity. Due to its hydrophobic character, it is plausible that HQNO directly interacts with the membrane to increase fluidity. In light of this result, we hypothesized that Antimycin A and Oligomycin, both hydrophobic compounds, also increase *S. aureus* sensitivity to chloroxylenol by altering membrane fluidity; the other ETC inhibitors tested, 3-NP and sodium azide, which did not enhance sensitivity to chloroxylenol, are both hydrophilic compounds. We showed that treatment of *S. aureus* with Antimycin A also leads to an increase in membrane fluidity. These findings suggest that the observed HQNO-mediated increase in antibiotic efficacy is independent of the effect of HQNO on the *S. aureus* ETC. Furthermore, we showed that modulating membrane fluidity via either genetic or chemical approaches shifts *S. aureus* chloroxylenol sensitivity profiles. Together, these results are consistent with a model whereby HQNO increases *S. aureus* membrane fluidity, which greatly enhances the ability of chloroxylenol to kill *S. aureus* biofilms.

We also found that treatment with *P. aeruginosa* supernatant or pure HQNO influenced the membrane fatty acid composition of *S. aureus*. Specifically, *S. aureus* grown in medium alone had a significantly higher proportion of BCFA compared to *S. aureus* cells exposed to *P. aeruginosa* supernatant or HQNO for 24 h. Given these results, we hypothesize that HQNO-mediated inhibition of the *S. aureus* ETC leads to decreased rates of fatty acid synthesis. Previous work from our laboratory has shown that when these organisms are in co-culture, *P. aeruginosa* forces *S. aureus* to grow by fermentation (84), which leads to a reduction in growth of *S. aureus* (37). Furthermore, during co-culture with *P. aeruginosa, S. aureus* downregulates multiple genes involved in fatty acid synthesis, including the cardiolipin synthase (*cls1*), and branched-chain amino acid transporters (*brnQ1, brnQ2, brnQ3, bcaP*) (84). Additionally, it has been shown that anaerobically-grown *S. aureus* has lower protein synthesis rates for multiple enzymes involved in metabolism, including FabG1, which is required for fatty acid synthesis (85).

Together, our results are consistent with the following two models, which are not mutually exclusive: 1) HQNO increases *S. aureus* membrane fluidity, potentially via direct interaction with the membrane, and 2) exposure to HQNO slows or halts *S. aureus* fatty acid synthesis, leading to altered membrane lipid composition, perhaps via ETC inhibition. Our data suggest that the first model may explain how HQNO potentiates the activity of chloroxylenol against *S. aureus* biofilms. In contrast, our data do not support a role for the second model in explaining the altered chloroxylenol susceptibility profiles we observe. Specifically, the HQNO-mediated increase in *S. aureus* membrane fluidity occurs more rapidly compared to the *P. aeruginosa* supernatant-induced changes in fatty acid profiles. Therefore, we hypothesize that HQNO increases fluidity via direct interaction with the membrane, rather than via inducing a shift in membrane fatty acid composition. The second model could explain other potential consequences of this interspecies interaction, such as an impaired ability to adapt to changing environmental conditions.

We observed that *P. aeruginosa* exoproducts can potentiate the activity of multiple membrane-active compounds, including the phenol biphenyl, and gramicidin, which forms channels within the membrane (86-88). Interestingly, we also showed that *P. aeruginosa*-secreted factors enhanced the activity of two non-traditional antibiotics, trifluoperazine and amitriptyline. Both of these drugs have a fused tricyclic structure, and have been found to possess antibacterial activity (89-93). Additionally, trifluoperazine was found to synergize with fluconazole against multiple fungal species (94). Due to its high degree of hydrophobicity, trifluoperazine has been shown to interact with cell membranes and cause increased fluidity and permeability (95, 96); it has been hypothesized that amitriptyline acts in a similar manner (93).

Importantly, we found that the combination of *P. aeruginosa* supernatant and chloroxylenol was effective against multiple slow-growing *S. aureus* populations, namely, anaerobically-grown biofilms and SCVs. From a therapeutic perspective, these results could have important implications. Infection sites can have steep oxygen gradients (97, 98), which may lead to slow microbial growth *in vivo* (99). Slow-growing pathogens are difficult to eradicate because many antibiotic classes are only effective against actively growing cells; in contrast, antibacterial agents that target membranes are effective whether or not bacteria are growing. Thus, our discovery that an interspecies interaction can potentiate the activity of membrane-active drugs could be used to inform the treatment of recalcitrant mixed-species infections involving bacterial biofilms in oxygen-depleted sites.

Overall, our work demonstrates that polymicrobial interactions can profoundly shift the antibiotic sensitivity profiles of bacteria growing as biofilms. In particular, we discovered that interspecies interactions can lead to changes in the fluidity and composition of the bacterial cell membrane, which may influence other aspects of bacterial physiology as well as responses to environmental stressors. Together, these findings may have important consequences for the treatment of polymicrobial infections in multiple disease contexts, including non-healing wounds and pulmonary infections in patients with cystic fibrosis. Finally, the knowledge we have gained from this work has the potential to inform the development of more effective anti-biofilm combination therapies.

## Materials and Methods

See the Supplemental Materials and Methods in Text S1 for additional details regarding the methods.

### Bacterial strains and culture conditions

A list of all strains used in this study is included in Table S2. *S. aureus* was grown in tryptic soy broth (TSB) and *P. aeruginosa* was grown in lysogeny broth (LB). All overnight cultures were grown shaking at 37°C for 12-14 h, except for the *S. aureus* Col *hemB* mutant, which was grown statically at 37°C for 20 h.

### Biolog MicroArray antibiotic susceptibility assay

Biolog Phenotype MicroArray bacterial chemical sensitivity assay panels were used to test *S. aureus* antimicrobial sensitivities as previously described (37). See the Supplemental Materials and Methods in Text S1 for additional details.

### Biofilm disruption assay on plastic

*S. aureus* biofilms were treated with antimicrobial agents, followed by enumeration of viable cell counts, as previously described (37). See the Supplemental Materials and Methods in Text S1 for additional details.

### Membrane potential measurements

*S. aureus* membrane potential was determined using the fluorescent dye DiOC_2_ as previously described (100, 101) with some modifications. See the Supplemental Materials and Methods in Text S1 for additional details.

### Laurdan membrane fluidity analysis

*S. aureus* membrane fluidity was determined by Laurdan generalized polarization (GP) as previously described (101, 102) with some modifications. See the Supplemental Materials and Methods in Text S1 for additional details.

### Fatty acid methyl ester analysis

Whole-cell direct fatty acid methyl ester (FAME) analysis of *S. aureus* pellets was performed by Microbial ID, Inc. (Newark, DE) as previously described (103). See the Supplemental Materials and Methods in Text S1 for additional details.

## Acknowledgements

We thank Ambrose Cheung, Deborah Hogan, Vineet Singh, and David Heinrichs for providing bacterial strains.

## Supplemental Figures

**Figure S1. The concentration of ethanol used in this study does not decrease *S. aureus* biofilm viability in the presence or absence of *P. aeruginosa* supernatant and *P. aeruginosa* supernatant rapidly increases *S. aureus* biofilm sensitivity to chloroxylenol.**

**Figure S2. Testing the ability of *P. aeruginosa* PA14 mutants defective in exoproduct production to increase *S. aureus* biofilm sensitivity to chloroxylenol.**

**Figure S3. Testing the ability of electron transport chain inhibitors and a proton ionophore to increase *S. aureus* biofilm sensitivity to chloroxylenol.**

**Figure S4. Reactive oxygen species do not sensitize *S. aureus* biofilms to chloroxylenol**

**Figure S5. Manipulating membrane fatty acid composition alters *S. aureus* biofilm sensitivity to chloroxylenol.**

**Figure S6. Exposure to *P. aeruginosa* exoproducts alters *S. aureus* membrane fatty acid composition.**

## Supplemental Tables

**Supplemental Table 1. *P. aeruginosa* supernatant increases *S. aureus* biofilm sensitivity to antiseptics and antibiotics.**

**Supplemental Table 2. Strains used in this study.**

**Supplemental Table 3. HQNO alters *S. aureus* membrane fatty acid profiles.**

**Supplemental Text 1. Supplemental Results and Materials and Methods.**

